# The role of early acoustic experience in song discrimination

**DOI:** 10.1101/756445

**Authors:** Emily J. Hudson, Nicole Creanza, Daizaburo Shizuka

## Abstract

Oscine songbirds are an ideal system for investigating how early experience affects behavior. Young songbirds face a challenging task: how to recognize and selectively learn only their own species’ song, often during a time-limited window. Because birds are capable of hearing birdsong very early in life, early exposure to song could plausibly affect recognition of appropriate models; however, this idea conflicts with the traditional view that song learning occurs only after a bird leaves the nest. Thus, it remains unknown whether natural variation in acoustic exposure prior to song learning affects the template for recognition. In a population where sister species, golden-crowned and white-crowned sparrows, breed syntopically, we found that nestlings discriminate between heterospecific and conspecific song playbacks prior to the onset of song memorization. We then asked whether natural exposure to more frequent or louder heterospecific song explained any variation in golden-crowned nestling response to heterospecific song playbacks. We characterized the amount of each species’ song audible in golden-crowned sparrow nests and showed that even in a relatively small area, the ratio of heterospecific to conspecific song exposure varies widely. However, although many songbirds hear and respond to acoustic signals before fledging, golden-crowned sparrow nestlings that heard different amounts of heterospecific song did not behave differently in response to heterospecific playbacks. This study provides the first evidence that song discrimination at the onset of song learning is robust to the presence of closely related heterospecifics in nature, which may be an important adaptation in sympatry between potentially interbreeding taxa.

## Introduction

Juvenile experience can set the stage for behavior later in life. The effects of early sensory experience have been studied in many taxa in the context of mate choice^1–6^ as well as in non-mating contexts^7–9^. How evolution shapes the timing and selectivity of learning has received considerable theoretical attention^10–12^ and, in a few cases, has been demonstrated empirically with adult mate choice^13,14^. Learning is especially well studied in oscine songbirds, in which exposure to acoustic cues influences what songs a bird will later sing^15–17^. Young birds that are not exposed to conspecific song tend to develop abnormal songs^18,19^. It is critical that songbird fledglings quickly identify conspecific models; failure to learn conspecific song can limit a male’s chances of successfully attracting conspecific mates (e.g.^20,21^). Accordingly, songbirds exhibit selective song learning, e.g. by preferentially learning song from their own species, as demonstrated by decades of laboratory studies^22–24^. Furthermore, learning is often limited to a sensitive period, beginning shortly after fledging, in which young birds are able to rapidly memorize song syllables they hear and will later sing ^15–17^ Songs heard after this developmental window are not sung later ^18^, nor are songs exclusively heard by nestlings (i.e. before the sensitive period) ever produced later ^25^, suggesting that imitative male song learning does not occur in the nest (although see the discussion of embryonic learning below). Female learning is less well studied, but there is evidence that female songbirds also learn songs and song preferences ^26–28^. How young birds of both sexes accomplish selective learning, avoiding mistakenly learning from heterospecifics, is a crucial adaptation that is likely important to evolutionary processes such as sexual selection and cultural evolution, as well as reproductive isolation between closely related taxa. This is an especially critical task in populations where closely related taxa coexist during the breeding season.

In order for selective song learning to take place, young birds must be able to discriminate between conspecific and heterospecific songs. This is hypothesized to be accomplished using an ‘acoustic template’, a neural representation of a key feature or features of appropriate conspecific song, which is in place at the onset of the learning period^18,29^. Some experimental evidence, such as preferential learning or responsiveness to conspecific songs in young birds cross-fostered by heterospecifics^24,30^, suggests that acoustic templates at the onset of song learning are ‘innate’— i.e., genetically encoded and not influenced by early acoustic exposure^31^. However, recent work also shows that songbirds are capable of responding to and learning from acoustic experiences as nestlings^32,33^, or even as embryos inside eggs ^9,34–39^. The body of work testing for early song discrimination has shown that species recognition is detectable at a relatively early age, including in wild nestling birds^30,40–43^. Combining behavioral tests of species recognition in nestlings with measurements of natural variation in early song exposure allows us to investigate the extent to which young birds are sensitive to their acoustic environment while they are still in the nest. The critical question is whether early experience affects the ability of nestlings to recognize their own species—specifically, whether early exposure to heterospecific song may act in opposition to the well-documented tendency of young songbirds to preferentially respond to conspecific song. Alternatively, exposure to heterospecific song could improve a bird’s ability to discriminate against other species’ songs^44^. Intuitively, the task of discriminating against heterospecific song to avoid costly learning mistakes should be most important when sister species breed at the same site (and thus may risk hybridization). Is the behavior of young birds in such populations affected by the acoustic presence of their sister species, or has selection acted to mitigate exposure to heterospecific song (e.g. by favoring nestlings that ignore heterospecific song)? The first step in addressing this question is establishing the amount of heterospecific song, if any, that is heard by nestling birds in a natural context. The responses of nestlings that are exposed to relatively more heterospecific song in the nest could then be compared with those nestlings that hear relatively less heterospecific song.

To address this longstanding question of the role of early experience on species recognition, we focused on the golden-crowned sparrow (*Zonotrichia atricapilla*), a large sparrow found in western North America. The golden-crowned sparrow’s sister species, the white-crowned sparrow (Z. *leucophrys*), breeds across a wide swath of the United States and Canada and is a model organism for studying the timeline of avian vocal learning (summarized in Figure 1). In white-crowned sparrows, as in many bird species, song is known to play a key role in mate selection^45^, making the task of learning the correct song critical for males’ reproductive success (females of both species do not sing during the breeding season). Their close phylogenetic relationship with the well-studied white-crowned sparrow provides a solid basis for inferring some aspects of the golden-crowned sparrow song learning process; namely, that learning begins only after fledging, at ~ 10 days after hatching. Moreover, in many parts of their breeding range, these two species breed simultaneously in the same treeline habitat, making them an ideal species pair in which to study how the presence of closely related species affects learning. It has been noted that hybridization often occurs across large genetic distances in birds, with viable offspring produced from hybridizing parent species separated by tens of millions of years ^46,47^. Despite this potential for hybridization, the formation of hybrid pairs between golden-and white-crowned sparrows appears to be very rare (never seen at this site during fieldwork in June and July of 2013, 2015, and 2017, although two possible F1 hybrids have been described outside the breeding season in California^48,49^). Thus, despite the apparently ample opportunity to hybridize, a reproductive barrier clearly exists between these two closely related taxa, although whether this barrier is based on pre-mating behavioral barriers is not known.

**Figure 1.**
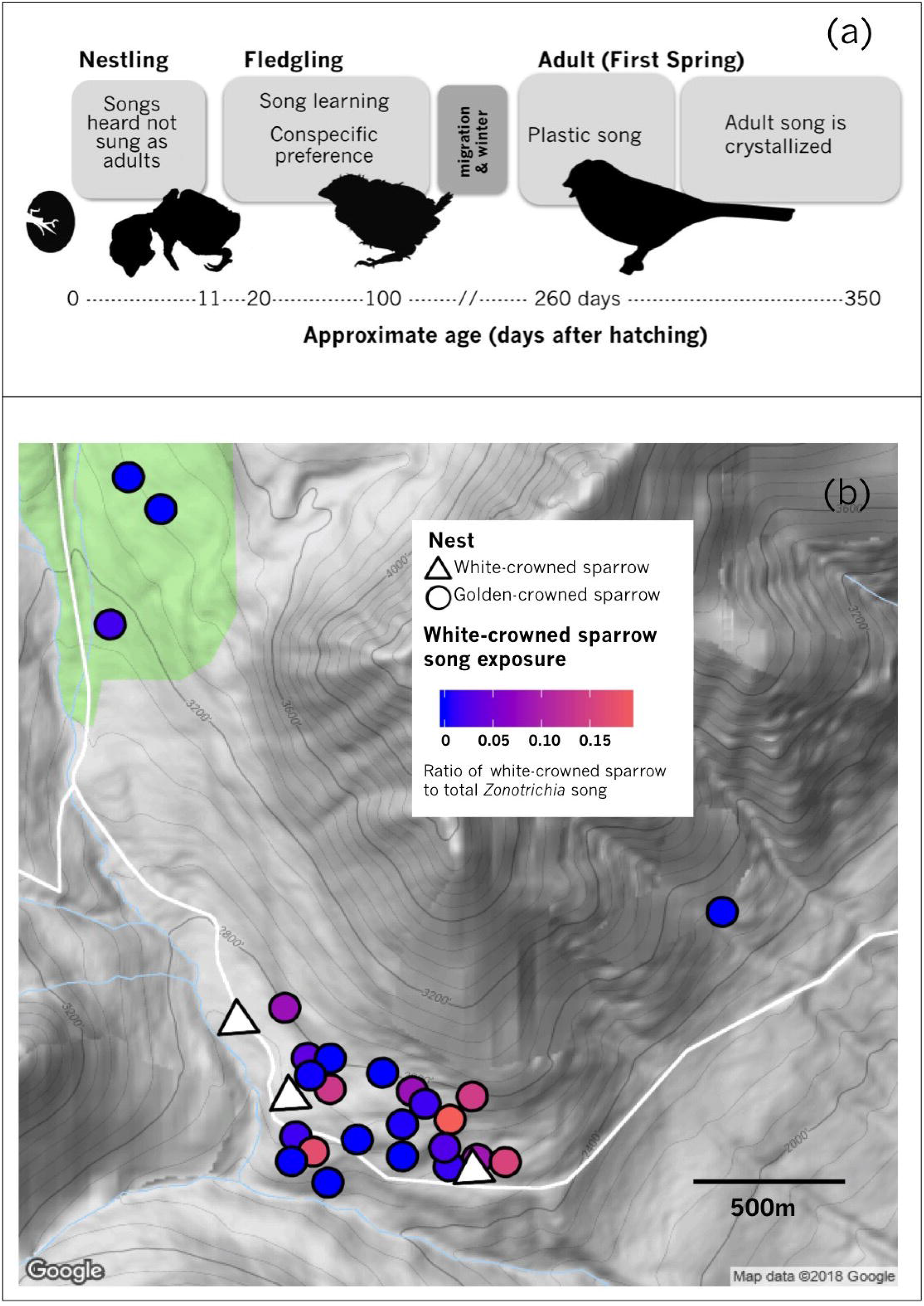
(**a**) Summarized timeline of song learning in white-crowned sparrows. The current study focuses on measuring the effect of experience in the closely-related golden-crowned sparrow at the nestling stage (day 8-10); at this age (prior to fledging at day 10-11;^79^), song learning is not thought to have begun^80,81^. (**b**) Positions of nests of playback subjects at Hatcher Pass Management area. White triangles represent active white-crowned sparrow nests; circles represent golden-crowned sparrow nests, with color representing the ratio of white-crowned sparrow song to total *Zonotrichia* songs recorded at that nest.

In this study, we take advantage of natural variation in heterospecific (white-crowned sparrow) abundance at our field site to test the effect of exposure to heterospecific signals on the behavior of nestling golden-crowned sparrows. We use golden-crowned sparrows to investigate (1) whether nestling golden-crowned sparrows can distinguish heterospecific and conspecific songs prior to the sensitive period for vocal learning, and (2) whether the acoustic environment in the nest affects this ability. If so, we might expect birds who hear more heterospecific song to respond differently to heterospecific stimuli than birds with less exposure to heterospecifics. Another possibility is that nestling birds only attend to the loudest songs audible at the nest, or those above some threshold amplitude; in this case, we would expect nestlings to respond most to the song types that are loudest at their nest. If nestlings show no early effect of acoustic experience, this would suggest that species recognition cannot be overwritten by heterospecific exposure, even when this exposure occurs at a very early stage.

## Methods

### Acoustic monitoring

All recordings and playbacks were conducted at Hatcher Pass Management Area, Alaska in June and July 2017. During this period, sunrise time varied between 4:11 and 4:46 a.m. Therefore, to capture the dawn chorus with a conservative margin of time before and after sunrise, we continuously recorded between midnight and 7 a.m. local time using five Wildlife Acoustics Song Meters (SM4)^50^. Nests were all constructed on the ground in this population, so recorders were placed on the ground within 1m of each golden-crowned sparrow nest for one dawn chorus per nest, within 24 hours of conducting playback experiments (N=23 nests, of which 19 were successfully used in playback experiments and retained for subsequent analysis). Recordings were saved as consecutive 20 minute WAV files, sampled at 16 or 24kHz (16 bits per sample). Six researchers annotated recordings, noting golden-crowned and white-crowned sparrow songs if they were visible on the spectrogram in the program Syrinx (J. Burt, Seattle, WA). Overlapping songs from different birds were counted as two separate songs if approximately 33% or less of the songs overlapped. White-crowned sparrow exposure was then calculated for each nest by dividing the number of white-crowned songs detected by the total number of golden-crowned and white-crowned songs detected.

We also quantified the relative amplitude of both golden-crowned and white-crowned sparrow songs. Relative amplitudes were compared only between songs recorded on the same day with the same SM4 unit. We generated a Gaussian-windowed spectrogram of these songs in Matlab (as in ^51^), which results in a time-by-frequency matrix of signal intensity. For each annotated song, we calculated mean and total amplitude of each song using the spectrogram matrix values within the time and frequency bounds identified in Syrinx. However, different nests had different levels of environmental background noise, which affects our amplitude calculations. To remove environmental background noise from our song amplitude measurements, we took the average amplitude from a period of time (between 12 a.m. and 1:30 a. m.) in the same recording without birdsong, and subtracted this average background value from the average amplitude of annotated songs for each nest (see Supplemental Fig 1 for an example). This provided an estimate of the relative amplitude of golden-crowned sparrow songs and white-crowned sparrow songs separately at each nest.

### Playback Experiment

Previous studies showed that nestling golden-crowned sparrows that were played either conspecific song or that of the sympatric white-crowned sparrow chirp more in response to conspecific songs^40,42,43^. To test the role of early song exposure in modulating this behavior, we followed the same protocol as previous studies by conducting playbacks when nestlings were about 8–10 days old, as estimated based on hatch date or length of exposed primary feather (>6mm for the majority of nestlings). All nestlings in a nest (2–6 per nest, mean 4.2) were temporarily removed and randomly assigned to one of two playback treatments (golden-crowned or white-crowned sparrow song), each consisting of 6 stimulus files created from a unique recording of a different individual male to avoid pseudoreplication (recorded >6 years prior at sites >100km away). These golden-crowned sparrow song recordings are of the same dialect type that males at this study site produce ^52^, and have been effective at eliciting strong responses from adult males and nestlings in this population previously ^42,43^. As in ^40,43^, each stimulus file consisted of one minute of white noise, two minutes of song presentation (the same song recording repeated every 10s), and an additional minute of white noise. For each trial, an individual nestling subject was placed alone in a collapsible cloth pet carrier (26×27×48 cm). Songs were broadcast from an iPod Nano mp3 player (Apple) through a speaker (SDI Technologies, Inc., Rahway, NJ) placed immediately outside the pet carrier. Playback volume was standardized to 60 dB at 1m from the speaker for consistency with previous studies in this population ^42,43,53^. The observer recorded the number of chirps the nestling produced during the 1 minute pre-playback period, 2 minutes of song playback, and 1 minute post-playback period. A previous study using the same protocol found high inter-observer agreement in chirp numbers when trial videos were re-scored later by a different individual^43^.

### Molecular Sexing

We collected nestling blood from the brachial vein immediately following playback trials and stored the blood on FTA filter paper cards. We determined the sex of individual nestlings using a standard DNA-based sexing protocol^54^, which has been validated for this species^55^.

### Statistical analysis

All statistical analyses were carried out using R (version 3.5.0, R Core Team 2018). We first used a linear model to measure the effect of white-crowned song exposure on responsiveness to white-crowned song by using the mean response to white-crowned sparrow song in each nest (which was heard by 1-3 chicks per nest) as the response variable. Because many individual factors can affect a nestling’s likelihood to respond, we also ran a generalized linear mixed-effects model with Quasi-Poisson regression, with individual responses as the response variable and several fixed effects: individual subject factors (pre-playback behavior, sex, and exposed primary length), nest-level factors (white-crowned sparrow song exposure, clutch size), and playback type (white-crowned or golden-crowned sparrow). We then used R package MuMIn^56^ to perform model selection based on ΔQAICc < 4. The retained models were averaged, and summaries of coefficients for each effect are presented in Table 1.

## Results

Nestling golden-crowned sparrows were able to discriminate behaviorally between conspecific and white-crowned sparrow song (Fig 2), supporting previous findings in this population ^42,53^. We found considerable variation in the amount of white-crowned sparrow song recorded at each golden-crowned sparrow nest. While some golden-crowned sparrow nests had no audible white-crowned sparrow songs (N=6), the majority of golden-crowned sparrow nests were exposed to some white-crowned sparrow song (Fig 2), at levels between 0.03–19% of the combined amount of golden-crowned and white-crowned-sparrow song (N=13). We found that white-crowned sparrows heard at golden-crowned sparrow nests were louder on average than golden-crowned sparrow songs (Wilcoxon rank sum test, *P* < 0.001, Fig 3a). We also found, from a three nests at which 24 hour recordings were annotated, that song production of both species was highest between 3 and 7 a.m. (Supp Fig 1).

**Figure 2.**
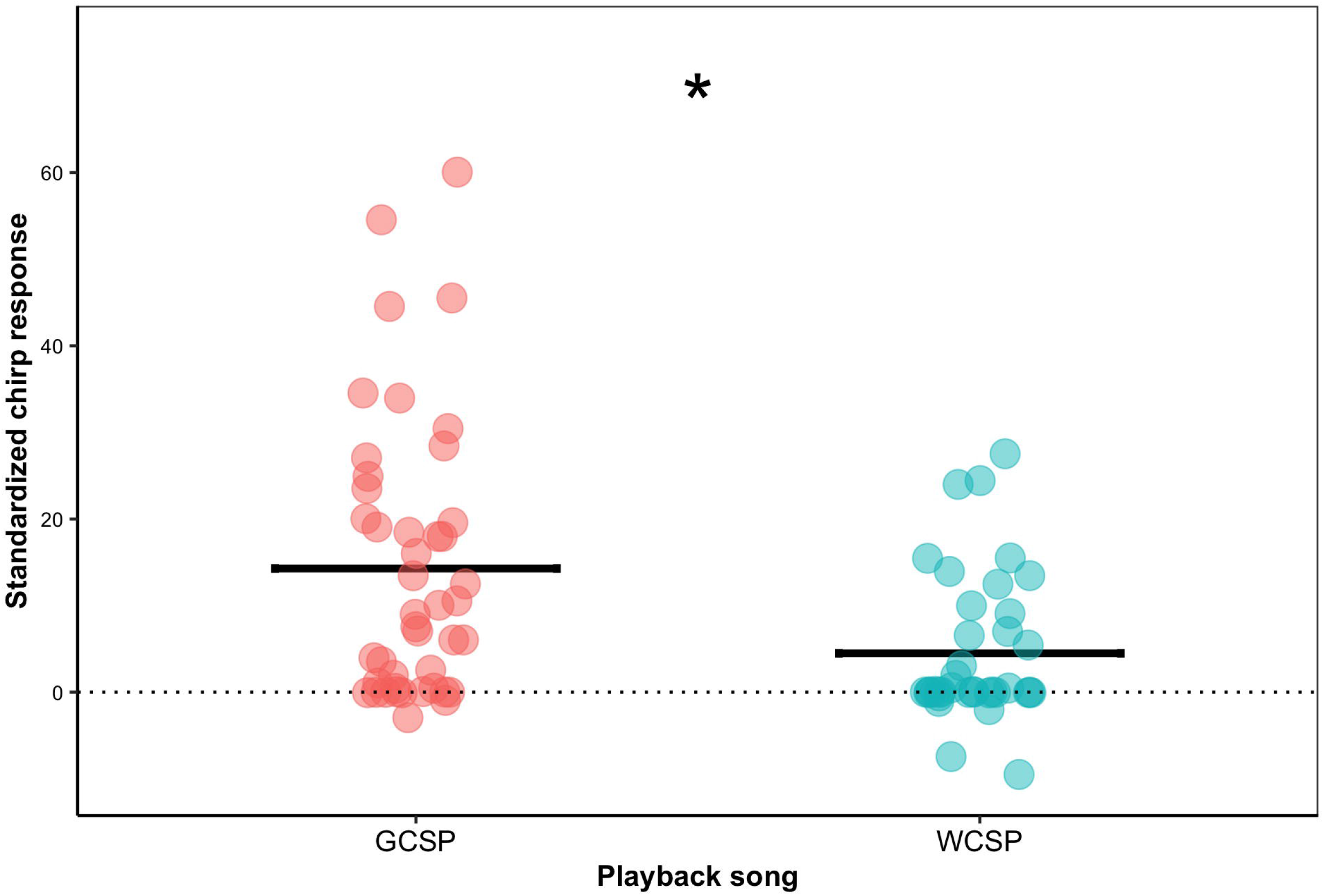
Response of golden-crowned sparrow nestlings to playbacks of conspecific (left) and white-crowned sparrow (right) songs. To better visualize responses, here the number of chirps in the one-minute pre-playback period was doubled and subtracted from the number of chirps during the two-minute playback period. Statistically, the effect of pre-trial chirping was accounted for by adding it as a fixed effect to the mixed-effects model (see Methods). The nestling response to conspecific, golden-crowned sparrow songs was significantly higher than to white-crowned sparrow songs (*P* = 0.0007).

**Figure 3.**
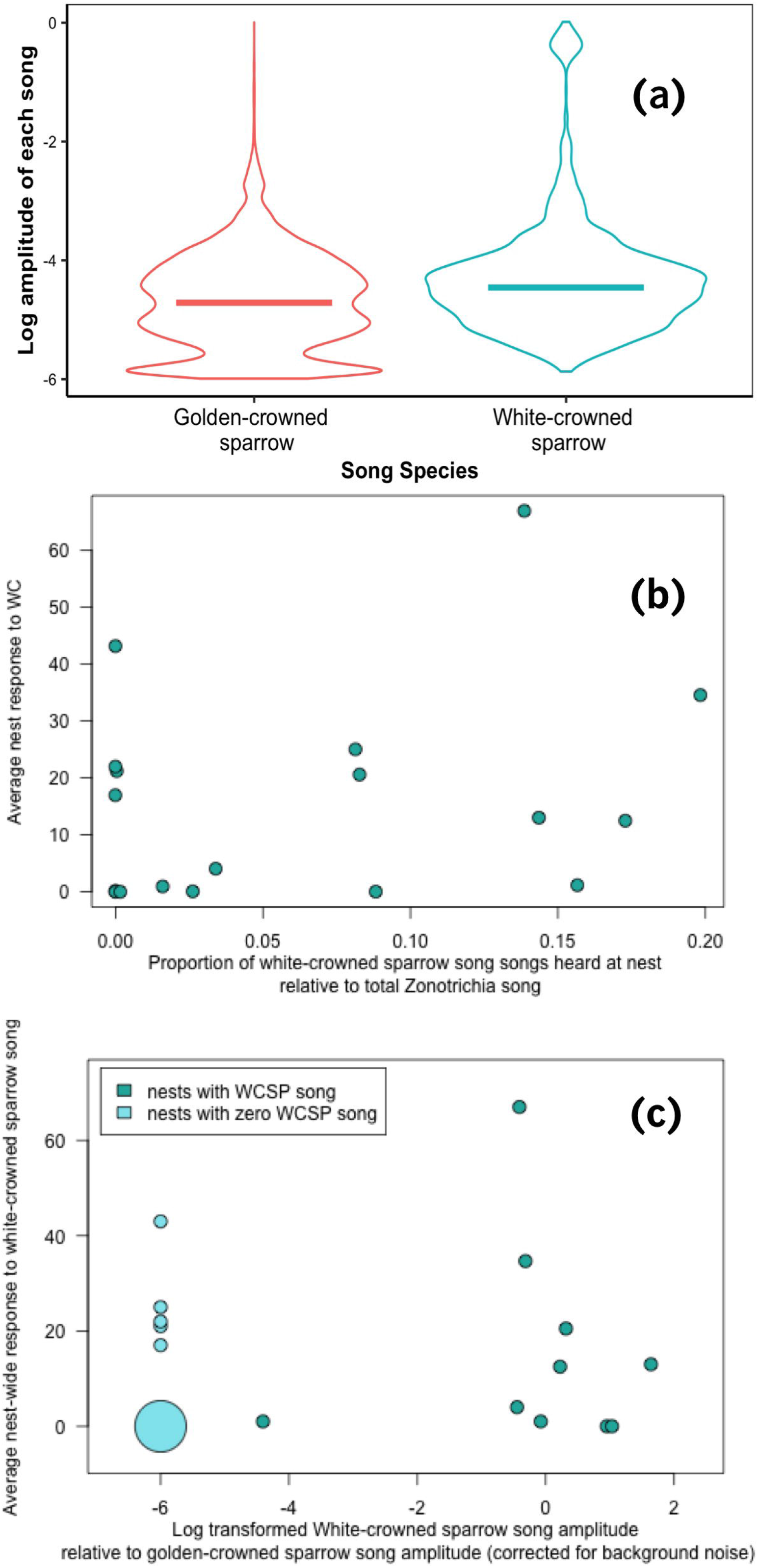
(**a**). The amplitudes of all recorded *Zonotrichia* songs at each golden-crowned sparrow nest (N=15), log-transformed to aid visualization. White-crowned sparrow songs, although less frequently heard, were on average louder than golden-crowned sparrow songs heard at golden-crowned sparrow nests (medians shown with horizontal bars; Wilcoxon rank-sum test, p<0.001). (**b**) Average response of golden-crowned sparrow nestlings to white-crowned sparrow playback, as explained by proportion of white-crowned sparrow song to total *Zonotrichia* song heard at the nest during the dawn chorus. Nest-wide averages were computed for the response variable for all nests that had more than one nestling receive the white-crowned sparrow treatment; if only a single chick received this treatment, that subject’s response was plotted. There was no effect of proportion of white-crowned sparrow song on average responsiveness. (**c**) Average response of golden-crowned sparrow nestlings to white-crowned sparrow song playback, as explained by relative amplitude of white-crowned sparrow songs at each nest. As before, nest-wide averages were computed for the response variable for all nests in which more than 1 nestling receive the white-crowned sparrow treatment. For each nest, a background noise level was computed from portions of the nightly recording without birdsong, which was then used to filter out songs of both species which were lower than that threshold. Ratios of average white-crowned sparrow amplitude to average golden-crowned sparrow amplitude are transformed to a log scale to aid visualization; larger negative values represent nests with higher ratios of golden-crowned sparrow amplitude relative to white-crowned sparrow amplitude, and *vice versa.* Nests with no white-crowned sparrow songs (zero amplitude) are represented in light gray and assigned to an arbitrary value of −1 on this scale. The radius of each circle represents the number of points at identical coordinates. There was no effect of relative amplitude of white-crowned sparrow song on average responsiveness.

Exposure to white-crowned sparrow song did not increase average responsiveness to white-crowned song at a given nest (linear model; *P* = 0.27, Figure 3b), nor did the amplitude of white-crowned song predict the response to white-crowned song (linear model *P* = 0.44, Figure 3c). Using the difference between average nest-wide chirps to golden-crowned and white-crowned sparrow song as the response variable produced similar non-significant results (*P* = 0.39), as did the proportion of average nest-wide chirp responses to white-crowned song over chirps to both song types (*P* = 0.32); therefore, we only present total individual chirp values hereafter to maintain consistency with previous studies in this population^40,42^. To account for individual differences in nestlings that we know affect likelihood to respond (e.g. differences in exposed feather length or number of chirps pre-stimulus)^40,42^, we ran a global linear model and used AICc model selection to identify the most important factors in variation in response. The top ranking models (ΔQAICc <4) all retained pre-track response, clutch size, white-crowned sparrow exposure, and an interaction between white-crowned sparrow exposure and playback type, but only some included sex. Overall, the number of chirps prior to playback, and the species of song playback (golden-crowned sparrow or white-crowned sparrow) had the greatest effect on golden-crowned nestling response: chicks tested with golden-crowned sparrow song, and chicks that chirped more during the pre-stimulus period, were likely to chirp more during the playback period. Interestingly, clutch size had a small but significant effect on chirp number (*P* = 0.042, Table 1 and Supplemental Figure 2) while white-crowned sparrow song exposure did not (*P* = 0.12, Table 1).

## Discussion

In this study of golden-crowned sparrows, we quantified the amount of conspecific song, as well as the amount of song of a congeneric sister species, audible in the nest in the wild. We show that the amount and amplitude of white-crowned sparrow song heard by golden-crowned sparrow nestlings does not influence their response to playbacks of either species’ song. Documenting the opportunity for exposure to heterospecifics, especially closely related species, is important for understanding the evolution of recognition and its role in reproductive isolation. We found that even within a small area (~0.5 km^2^), nests of golden-crowned sparrows varied considerably in their amount of white-crowned sparrow song exposure. Interestingly, the level of exposure did not seem to be purely explained by distance to the closest white-crowned sparrow nest (Figure 1). This may be explained by the natural history of territoriality and singing behavior in these species: in both species, males repeatedly use singing perches that are within their territory but not necessarily close to the nest (pers. obs.). In species pairs that have overlapping territories, this can lead to the counterintuitive pattern that the heterospecific neighbors’ songs can be heard louder at a nest than the songs of the father or conspecific neighbors. We are confident that the nests with high levels of white-crowned exposure in the core area of the study site are not explained by the presence of undetected white-crowned sparrow nests nearby; since all male birds on our study site were banded, an unbanded male white-crowned sparrow with a nest in this area would almost certainly have been noted during our ~8 hours of daily observations. Our findings suggest that even within the same population, individuals likely experience widely varying acoustic environments. This intuitive but rarely documented fact should be considered when studying learning, discrimination, and sympatry in the wild.

The present study is consistent with previous results suggesting that conspecific songs are more salient to nestling birds than heterospecific songs^40,42^. This has often been explained as an innate predisposition, consistent with results from young birds kept in isolation. An intuitive alternative explanation is that this predisposition is shaped by very early acoustic experience. We tested this alternative hypothesis here, and found no nest-wide effect of ambient heterospecific song exposure on golden-crowned sparrow nestling responses to heterospecific playbacks; on both an individual and nest-by-nest basis, white-crowned sparrow song exposure did not explain a significant amount of variation in nestling response. Only the playback type (golden-crowned sparrow or white-crowned sparrow song) and pre-playback activity level (chirps prior to the start of the stimulus) predicted response to the playback. Thus, our results support a commonly assumed^57^, but difficult to test, idea that preferential responses towards conspecific songs in fledglings are not learned via experience in the nest.

Many songbirds, including sparrows in the genus *Zonotrichia,* show a large degree of within-species geographic variation in their songs^52,58,59^; our results raise new questions about how early experience with varied dialects affects behavior in young birds. Nestling golden-crowned sparrows in the present population discriminate against foreign dialects of conspecific song, responding to playbacks of these unfamiliar dialects as little as to white-crowned sparrow song^43^. This suggests that nestlings either have an innately determined preference for local song, or, in contrast to our results with heterospecific song here, nestling experience with local conspecific song increases its salience. In some taxa^14,60^, selection is hypothesized to favor genetic assimilation of learned responses (e.g. innately encoding the ability to discriminate against heterospecifics) when learning errors are costly. It may be the case that in golden-crowned sparrows, learning from white-crowned sparrows is so costly that nestlings have evolved to filter out their songs as nestlings, setting the stage for accurate learning later in development. However, conspecific stimuli may still pass this early filter, and lead nestlings to respond preferentially to the local dialect they hear most often. This explanation fits our results and is consistent with the theory that young birds might initially accept only a narrow range of conspecific signals based on filial imprinting, which is later broadened via experience with other conspecifics^61^. Manipulating nestlings’ acoustic environment to include foreign variations of conspecific song, as in ^62^, prior to nestling playbacks would further clarify the effects of early experience.

Although clutches at this site almost always consist of 4 or 5 chicks, interestingly, in this experiment the 2 largest clutches, with 6 chicks each, chirped more than average to all stimuli, such that clutch size was a significant fixed effect in our model comparisons. This result should be interpreted cautiously, since so few nests seem to be driving this pattern. However, behavioral (increased begging ^63^) and physiological (increased testosterone^64^ and sometimes corticosterone ^65,66^) effects of large clutch size have been documented in other songbirds where brood size was manipulated. One possible connection between clutch size and vocal response may be that chicks in larger broods are hungrier, or must vocalize more to solicit the same amount of parental provisioning, relative to chicks in smaller broods. We note that in the present study, the chirp response we measured corresponds to what Marler called the “fledgling location chirp”^22^, rather than a begging call, and is quite distinct from the broadband begging vocalizations chicks make in the nest during feeding. Therefore, we consider it unlikely that there is a straightforward relationship between hunger and chirping as measured here. Nevertheless, developmental stress caused by larger brood size (and independent of hunger) may still manifest itself in behavioral responses. Developmental stress has been shown to affect song learning by reducing copying accuracy when brood size is increased^67^, and reducing learned song complexity when corticosterone is administered^68^. Whether increased nestling response to playbacks is related to impaired learning later in development warrants further investigation.

Historically, selective song learning has been thought to begin only after fledging, as adult male white-crowned sparrows do not sing conspecific songs that were presented exclusively during the nestling period^22,25^. However, it is possible for birds to learn earlier in development than the fledgling stage: nestlings have been shown to learn familial contact calls^69^ and brood parasites can adapt their begging calls to their host species^32,33,70^. In addition, embryonic birds of multiple species have been shown to respond to ^71,72^ and even learn from ^9,34,39^ species-specific sounds. Therefore it is crucial to rigorously test for early learning, which has been difficult in songbirds for two main reasons. First, a lack of song production does not necessarily mean a lack of learning: birds may retain songs in memory^73^, even if they do not produce them as part of their adult repertoire. Secondly, the most straightforward test of the role of very early learning requires raising birds in acoustic and social isolation from hatching (or even earlier, to avoid embryonic learning^34^), which is a logistical challenge in most songbirds.

An alternative approach is cross-fostering, in which nestling birds are raised by a heterospecific foster parent, to see what effect early exposure to heterospecifics has on later behavior^74,75^. These studies generally measure mate choice in adults, and so cannot directly test when in development heterospecific experience is most influential. We assayed behavioral discrimination at the nestling stage to test for early learning in real time. Our findings are consistent with the standard model of song learning, in which nestlings possess an innate, species-specific auditory template that is not overwritten by early experience.

To better understand how recognition systems can evolve, this study focused on measuring naturally occurring variation in heterospecific song exposure in a population dominated by conspecifics. However, artificially manipulating the proportion of heterospecific song exposure to include more extreme values would also be informative. Perhaps heterospecific exposure below some threshold does not influence juvenile behavior, but at high levels, recognition is affected. Manipulating nestlings’ experience by broadcasting high levels of heterospecific song at the nesting site, similar to the approach used by Mennill *et al*.^62^, would be an important step in understanding learned recognition when abundances are unequal, such as at range limits or in hybrid zones.

Learning—the ability to modify behavioral preferences or signals based on early experience—provides a non-genetic pathway for trait inheritance^10,21,76^. How learning mechanisms themselves evolve, and how cognitive constraints on learning in turn affect species interactions and reproductive isolation, is an important but difficult problem in evolutionary biology. In particular, when critical behaviors must be learned, the potential for costly learning mistakes arises; we would expect that selection acts to balance the benefits and costs of flexibility by favoring inherited mechanisms that minimize the likelihood of mistakes. These mechanisms should be in place prior to learning, a prediction that has traditionally been very difficult to test. Early exposure to heterospecific sounds in the nest could potentially affect preference in two ways: either it could enable more effective species recognition, strengthening a predisposition to respond to conspecifics^44^; or it could interfere with species recognition and diminish the difference in response to heterospecifics *versus* conspecifics^77,78^. In contrast, our finding—that a significantly lower response to heterospecific song playback was maintained despite wide natural variation in heterospecific song exposure in the nest—provides a new line of support for the prediction that early-life experience with heterospecifics is not enough to override species-selective learning mechanisms. Disentangling the role of genetically inherited predispositions *versus* experience requires insight into the details of the learning process, natural history, and the ecological context in which learning takes place. Due to their history in behavioral ecology research and their unique biogeography, white-and golden-crowned sparrows are a promising system in which to study the interaction of these factors.

