## Supplemental Figures for "The role of early acoustic experience in song discrimination"

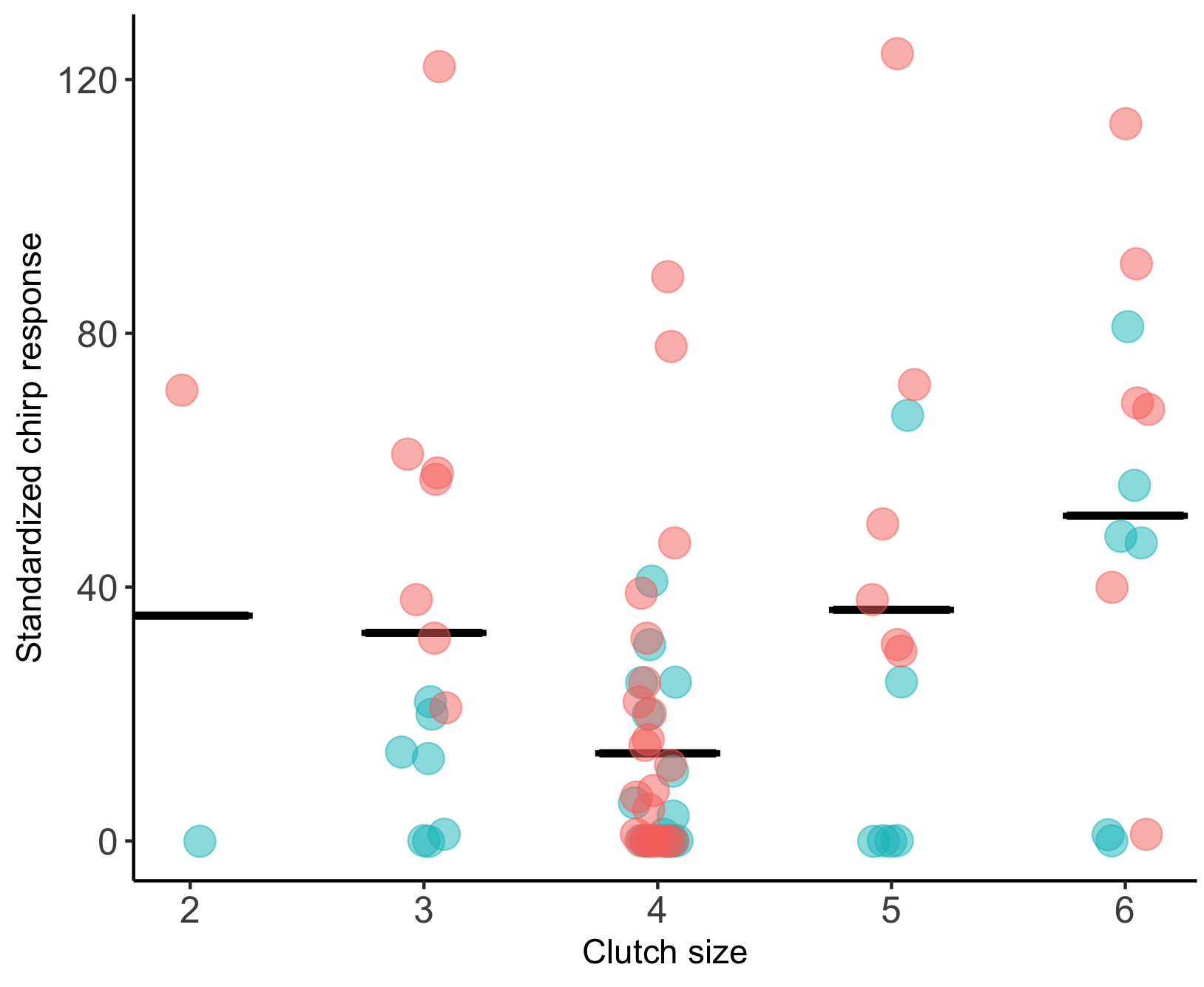
**Supplemental Figure 1.** Clutch size significantly affected nestlings’ likelihood to respond to playbacks, independent of which species song was played. Dots represent an individual golden-crowned sparrow chick’s response (number of chirps) to a playback (blue = white-crowned sparrow playback, red = golden-crowned sparrow playback).

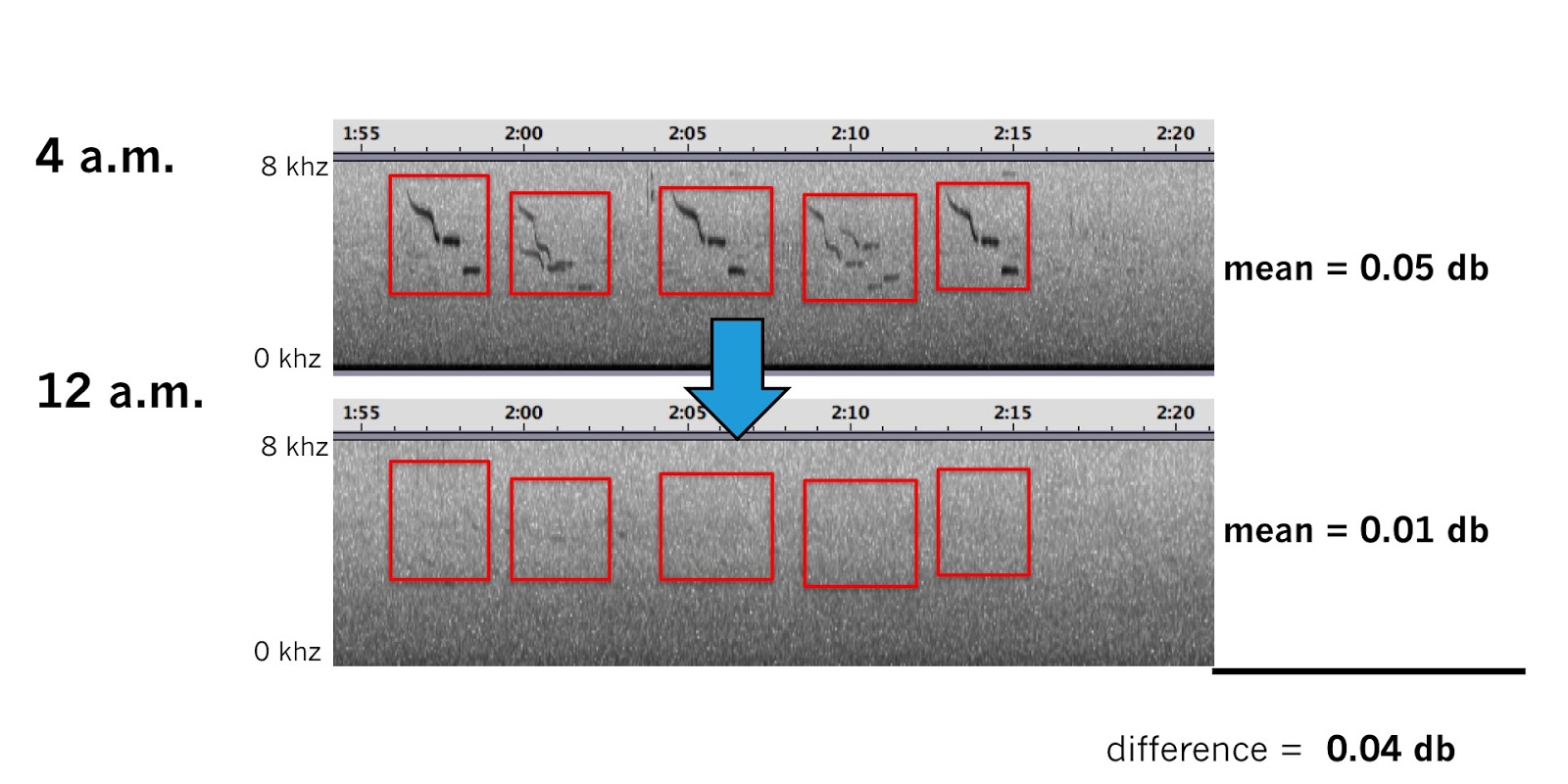

**Supplemental Figure 2.** A short example of how background noise amplitude was calculated for each nest. Songs were annotated for all continuous recordings (12 a.m. - 7 a.m.) at a single nest, and amplitudes within each box were calculated as described in Methods. To obtain the level of background noise (a value unique to each nest, in large part determined by distance to a nearby stream) we then took the 20 minute recording with the *most* songs in it (usually at or just before sunrise) recorded at that nest, and ran the same amplitude calculating procedure as before, but applying the annotations to the 20-minute recording with the *fewest* songs (shown in the example above as the 12 a.m. recording). We then subtracted the average amplitude from the boxes without songs in them from the average amplitude total for the nest, to estimate the amplitude of songs minus the environmental background noise.

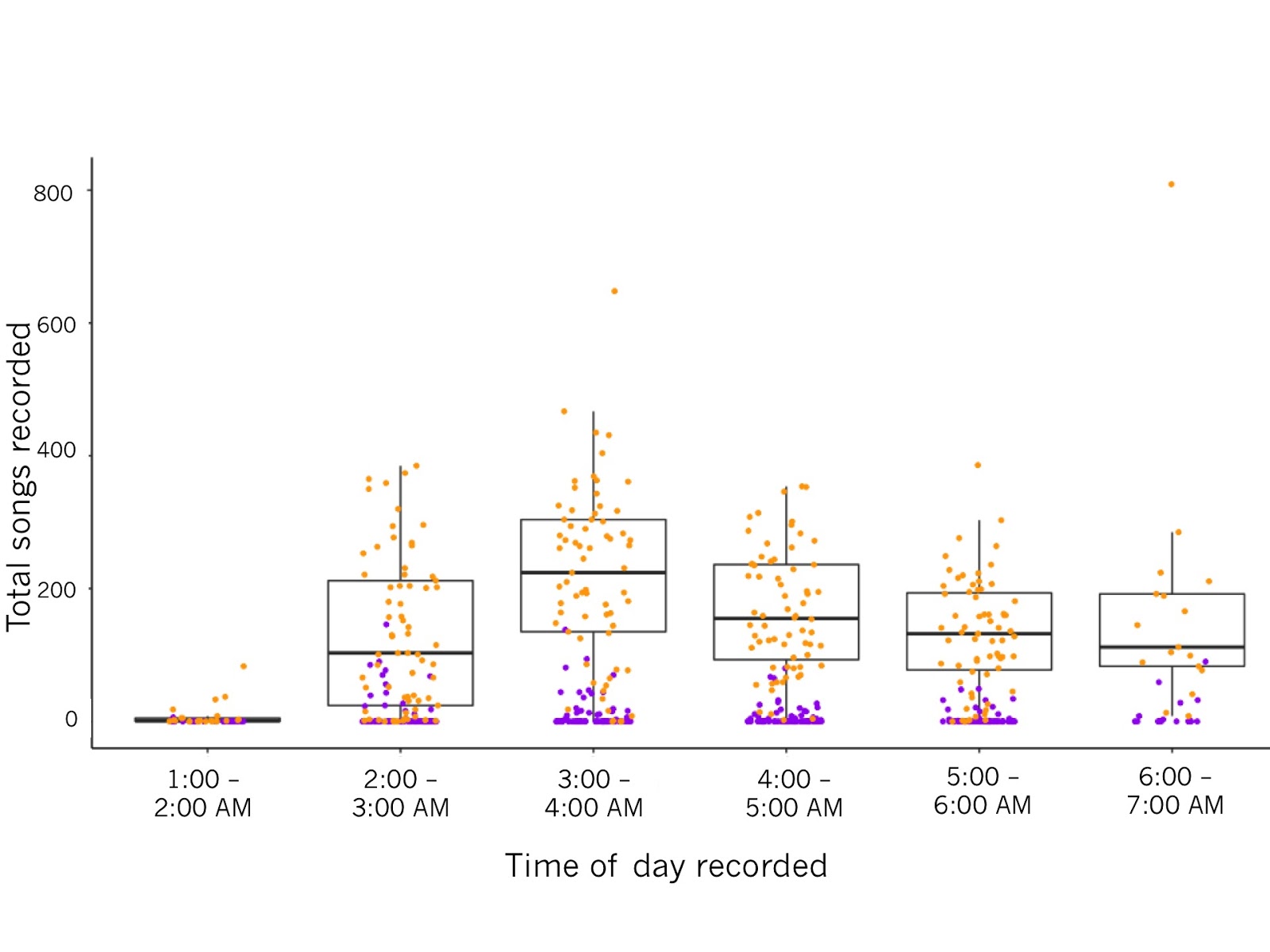

**Supplemental Figure 3.** Total songs counted in all nest (N=23) recordings for each species, binned by hour. Each dot represents the songs from a 20-minute sound file made during continuous recording. Box plots and orange dots represent golden-crowned sparrow songs, and purple dots represent white-crowned sparrow songs. Lines in the center of each boxplot represent the median number of songs recorded for that hour.
